# Are mosquito species present in Spain competent for Oropouche virus?

**DOI:** 10.64898/2026.01.23.701265

**Authors:** Rafael Gutiérrez-López, Nuria Labiod, Marcos López-de-Felipe, Patricia Sánchez-Mora, Sarah Delacour-Estrella, Alexandra Martín-Ramirez, Arantxa Potente, Eva Pérez-Martínez, Ricardo Molina, Maripaz Sánchez-Seco, Maribel Jiménez, Ana Vázquez, Inés Martín-Martín

## Abstract

Oropouche virus (OROV; *Orthobunyavirus*) is an emerging arbovirus endemic to South America and the Caribbean, with imported cases in European countries, including Spain. Although primarily transmitted by biting midges (*Culicoides* spp.), OROV has been detected in several mosquito species, raising concerns about potential establishment in non-endemic regions. European populations of *Aedes albopictus*, and *Culex pipiens*, as well as the invasive *Aedes aegypti*, represent relevant models for assessing vector competence. Here, we evaluated vector competence of Spanish *Cx. pipiens* biotype *molestus*, Spanish *Ae. albopictus*, and *Ae. aegypti* (Liverpool strain), for the 2024 OROV outbreak strain. Female mosquitoes were orally exposed to infectious blood meals and maintained under controlled insectary conditions. In addition, an additional group received a second non-infectious blood meal. The survival of the mosquitoes was monitored, and infection, dissemination, and transmission rates were assessed at 7-, 14-, and 21-days post-infection. Vertical transmission of the virus to the progenies was also analyzed. Overall, *Ae. albopictus* exhibited low infection rates, with occasional dissemination and transmission events. *Aedes aegypti* and *Cx. pipiens* showed infection and occasional dissemination, but no evidence of transmission. A second non-infectious blood meal did not significantly affect infection, dissemination, or transmission rates in any species. Viral loads in bodies and legs were low and did not differ significantly between species, time points, or feeding regimens. Survival was not affected by infection or blood-feeding regime. We did not find vertical transmission of OROV to the progenies. Regardless of virus dissemination in mosquitoes, our study indicates poor vector competence of Spanish *Ae. albopictus* and a lack of competence in *Ae. aegypti* and *Cx. pipiens* for the circulating OROV strain. These findings suggest a low risk for local OROV establishment in Spain, although continuous surveillance and research are warranted to monitor potential vector-virus adaptation.

## Background

Oropouche virus (OROV; *Orthobunyavirus*) is an emerging and neglected arbovirus endemic to South America and the Caribbean. However, in 2024, an epidemic of Oropouche fever resulted in more than 13,000 confirmed cases, including two deaths, reported in over 11 countries from America [1]. This outbreak has renewed attention to OROV as an important public health concern. Although no autochthonous cases have been notified in Europe to date, a total of 30 imported cases have been identified in travellers returning from endemic regions, with Spain reporting the majority of these cases (n = 21) [2, 3].

Current evidence indicates that OROV is primarily transmitted by biting midges of the genus *Culicoides* (Ceratopogonidae), particularly *Culicoides paraensis*, which is considered the main vector in urban cycles [4]. However, OROV has also been detected in several mosquito species [5–12]. The virus was first isolated from a pool of *Aedes serratus* in Brazil in 1961 [12], and *Culex quinquefasciatus* has been proposed as a secondary, urban, anthropophilic vector after multiple isolations [4–6, 11]. Despite these findings, infection rates in mosquitoes have generally been low, suggesting limited susceptibility. Indeed, experimental oral infection studies have shown that *Cx. quinquefasciatus* is unlikely a competent vector of OROV [6, 8, 11].

The introduction of OROV into non-endemic regions through viremic travellers raises important questions regarding the competence of local *Culicoides* and mosquito populations to transmit the virus. There is increasing interest in Europe in assessing the vector competence of *Culicoides* and mosquito species, and consequently several research groups across Europe have recently published data on the vector competence of local species in their respective countries [13–15]. Spain has reported the highest number of OROV cases in Europe [3], highlighting the relevance of assessing the vector competence of mosquito species present in the country. In this context, both the invasive Asian Tiger mosquito, *Aedes albopictus*, and the autochthonous common mosquito, *Cx. pipiens*, are present throughout the country, and could potentially be relevant models for OROV vector competence studies. Furthermore, *Ae. aegypti*, another potential model, while not present in mainland Spain, has been occasionally reported in the Canary Islands, Spain, in recent years [16, 17]*. Culex quinquefasciatus*, a member of the *Cx. pipiens* complex, is absent from Spain, but *Cx. pipiens* is widespread throughout Spain and could potentially act as a vector.

The unprecedented scale of the 2024 outbreak may be associated with a novel viral reassortant. This reassortant contains an M segment from strains detected in the eastern Amazon between 2009 and 2018, and L and S segments from strains circulating in Peru from 2008 to 2021 [18]. Such a reassortant replicates to higher titres in mammalian cells and shows reduced sensitivity to neutralization by human OROV immune sera collected before 2016 [19]. These genetic changes may influence viral pathogenicity and symptomatology [19]. Indeed, during the 2024 outbreak, OROV infection was linked to fatal cases in previously healthy young women, as well as to fetal death and microcephaly [20]. Nevertheless, the consequences of viral genetic variation for the mosquito-mediated transmission of OROV remain poorly understood. Evidence from other arbovirus–vector systems suggests that even subtle genetic changes can have major effects on vector competence. For example, in the case of Chikungunya virus (CHIKV), a single amino acid substitution was shown to increase viral adaptation to *Aedes albopictus*, facilitating epidemic spread in areas where *Aedes aegypti* was absent [21]. Still, given the emergence of a novel reassortant strain, changes in viral pathogenicity or vector range cannot be excluded.

Moreover, increasing evidences show that acquisition of a second, non-infectious blood meal can significantly enhance arbovirus dissemination and shorten the extrinsic incubation period in mosquitoes [22–24]. This phenomenon is thought to result from transient microperforations in the basal lamina surrounding the midgut, facilitating viral escape into the hemocoel and dissemination throughout different tissues [22–24]. Such effects have been documented for several virus–vector systems [22–24] and may also apply to OROV.

Here, we evaluate the vector competence of *Cx. pipiens* biotype *molestus*, and *Ae. albopictus* populations originally collected in Spain, as well as the *Ae. aegypti* Liverpool strain, for OROV. Additionally, we assess the capacity for vertical transmission of OROV and investigate whether a second non-infectious blood meal influences vector competence. This study aims to determine whether mosquito species present in Spain have the potential to support OROV transmission and thus represent an emerging risk for local virus establishment.

## Material and Methods

### Insects

An established colony of *Cx. pipiens* biotype *molestus* (originally collected from Arganda del Rey, Madrid, Spain, in 2024 and molecularly identified by PCR, targeting the CQ11 microsatellite region [25]), and an established colony of *Ae. albopictus* (collected in Barcelona, Spain in 2009) [26] were used during the assays. Additionally, an *Ae. aegypti* (Liverpool strain) laboratory colony was also included. All colonies were maintained at the National Center for Microbiology, Instituto de Salud Carlos III (Spain), under controlled insectary conditions (27 °C, 12:12 h light–dark photoperiod, 80 ± 5% RH) in an environmental chamber (Thermo Fisher Scientific, Waltham, MA, USA). Adults were provided with 30% sucrose solution *ad libitum* until infection. Mosquitoes were reared at the BSL-2 insectary and transferred to the BSL-3 insectary for vector competence studies.

### Virus and cells

The strain OROV SP2024 (GenBank ID: PQ329253) was used for experimental infections. This strain was originally isolated by the Laboratory of Arbovirus at the National Center for Microbiology, from a traveller returning from Cuba during the 2024 OROV outbreak [3]. Viral stocks were propagated once in Vero E6 cells (CRL 1586, American Type Culture Collection (ATCC), https://www.atcc.org), incubated at 37 °C for 4 days after which cytopathic effects were observed using a microscope, and freshly harvested supernatant was directly used for mosquito infections. The use of freshly propagated virus leads to higher infection and replication rates in mosquitoes compared with frozen aliquots, as has been demonstrated in other arboviruses, such as Zika virus (ZIKV) [27]. Viral titers were determined by tissue culture infectious dose 50 (TCD₅₀) assay on Vero E6 cells, following standard protocols to quantify infectious viral particles [28].

### Vector competence assay

Vector competence experiments were conducted with 3–7-days-old female mosquitoes. Approximately 100 females per species were placed in 450 ml cardboard cups covered with mesh custom-made by Alenta (https://www.alenta.org/), and were transferred to the BSL-3 insectary for mosquito infection and follow-up. Independent biological replicates were conducted for *Ae. albopictus* (n=3), *Ae. aegypti* (n=2), and *Cx. pipiens* (n=5). Mosquitoes were deprived of sugar 24 h prior to feeding to stimulate blood ingestion. The infectious blood meal was prepared as a mixture of 2 parts of blood and 1 part of virus in a final volume of 3 mL and consisted of freshly collected defibrinated rabbit blood provided by a rabbit farm (San Bernardo Farm, Navarra, Spain) spiked with fresh OROV to a final titre of 2.5 × 10^7^ focus-forming units [TCD50]/ml, and supplemented with 5 mM ATP and 2.5% saturated sugar solution to stimulate blood-feeding. Mosquitoes were exposed to the infectious mixture through quail skin as a membrane at 37 °C for 45 min using a Hemotek feeding system (SP6W1-3; Hemotek Ltd, Blackburn, UK). After feeding, mosquitoes were anesthetized with CO2, and fully engorged females were separated on a chilled plate and transferred to new 450 ml cups and maintained with 30% sucrose *ad libitum* in an environmental chamber following the aforementioned rearing conditions at the BSL-3 insectary. Freshly engorged mosquitoes were collected at 0 days post infection (dpi) to confirm the acquisition of virus by RT-qPCR, indicating successful virus uptake. For each species and experimental condition, between two-four mosquitoes were collected and processed individually. A subset of engorged females was offered a second non-infectious blood meal 3 dpi consisting of fresh rabbit blood following the same feeding protocol. Those females that successfully took the second blood meal were assigned to the “double-blood-meal feeding group” (from here 2BF), whereas those that did not feed again were maintained as the “single-blood-meal feeding group” (from here 1BF). The mosquitoes from both groups were introduced in 450 ml cardboard cups and provided with 30% sucrose solution *ad libitum* and kept in the environmental chamber (Climacell®, MMM Group, Germany) at 27 °C, 80% relative humidity (RH) and 12:12 light: dark cycle, in the BSL-3 insectary. Survival was monitored daily from Monday to Friday up to 21 dpi for the 1BF group and 14 dpi for the 2BF group.

At 7, 14, and 21 dpi, mosquitoes were anesthetized with CO₂, legs were removed, and individuals were induced to salivate. For this, cold-anesthetized mosquitoes were immobilized onto autoclave adhesive tape by sticking their wings and bodies to the tape. Their proboscis was introduced into 10 µl pipette tips containing 10 µl of minimum essential medium (MEM supplemented with 50 µg/ml penicillin-streptomycin, 50 µg/ml gentamicin, and 2.5 µg/ml fungizone) and salivation was induced by topically applying 1 µl of 2% pilocarpine (Novartis 2012, Alcon Cusí S.A. Barcelona, Spain) on the thorax. After 30 min, the MEM–saliva mixture retained in the pipette tip was added to a tube containing 300 μL of MEM supplemented with 50 µg/ml penicillin-streptomycin, 50 µg/ml gentamicin, and 2.5 µg/ml fungizone. Each mosquito body and corresponding legs were placed in separate microtubes containing 300 µl of the same medium. Samples were manually homogenized with a sterile pellet pestle until complete tissue disruption was achieved, centrifuged at 6,000 g for 5 min, and supernatants were stored at –80 °C until analysis.

To evaluate vertical transmission of OROV, after the infectious blood meal, 24 *Ae. albopictus*, 5 *Cx. pipiens,* and 26 *Ae. aegypti* engorged female mosquitoes were allowed to lay eggs (first gonotrophic cycle, F1) into plastic cups partially filled with water in which filter papers were placed. The eggs were allowed to hatch, and the larvae were reared in the environmental chamber in the BSL3 and feed with TetraMin XL Flakes (Tetra, Germany) until adult emergence. For each species, five pools of five females were tested for possible transmission of the virus to the F1 generation.

All experimental infections and mosquito maintenance were conducted in a biosafety level 3 (BSL-3) insectary at the National Center for Microbiology, Instituto de Salud Carlos III (Spain), following institutional biosafety and biosecurity regulations.

### Forming assay

To determine vector competence status, we tested viral RNA in the body, legs and saliva samples of individual mosquitos by specific RT-qPCR for OROV detection [29]. All samples with a quantification cycle (Cq) value < 38 were considered positive [3, 29]. Infection rate (IR), dissemination rate (DIR) and, transmission rate (TR) were calculated as following: IR = proportion of mosquitoes with a positive body among the total number of mosquitoes exposed to the infectious blood meal that had been analyzed; DIR = proportion of mosquitoes with positive legs among mosquitoes with a positive body; TR = proportion of mosquitoes with positive saliva among mosquitoes with a positive body.

### Statistical analysis

Mosquito survival was analyzed using Kaplan–Meier survival analysis, and differences between individuals fed an infectious or a non-infectious blood-meal were assessed using log-rank (Mantel–Cox) tests. Data were grouped by species and feeding status (1BF, 2BF). Due to the low number of control samples, survival rates of 2BF *Cx. pipiens* are not shown. Survival among the evaluated species for 1BF was also assessed.

To evaluate IR and DIR, independent generalized linear mixed-effects models (GLMMs) with a binomial error distribution and logit link function were fitted. Mosquito species, blood-feeding groups (one or two blood meals), and days post-infection (7, 14 and 21 dpi) were included as fixed factors, with the variable of the experimental biological replicate included as a random effect to account for variation among independent assays. Interaction terms between species and blood-feeding regime were tested to evaluate species-specific effects of the second blood meal on the IR. Because only one saliva sample tested positive, statistical analysis for TR was not conducted due to insufficient statistical power. Viral load data (log₁₀-transformed RNA concentration) from body and leg samples were analyzed using linear mixed-effects models (LMMs) with the same fixed and random structure. In all the models, residuals were inspected for normality and homoscedasticity. No overdispersion was observed in all the models developed, supporting an adequate model fit. All analyses were performed in R (v4.2.3; R Core Team, 2025) using the lme4 package. All plots were generated using GraphPad Prism version 10.0.3 (GraphPad Software, San Diego, CA, USA). Statistical differences were considered for p <0.05.

## Results

A total of 1,044 *Aedes albopictus*, 295 *Aedes aegypti*, and 728 *Culex pipiens* females were exposed to an infectious blood meal. Feeding success differed markedly among species, with 228 (21.26%) *Ae. albopictus*, 250 (84.75%) *Ae. aegypti*, and 141 (19.37%) *Cx. pipiens* successfully engorged. Among engorged females, a second, non-infectious blood meal was taken by 54/134 (40.30%) *Ae. albopictus*, 35/122 (28.69%) *Ae. aegypti*, and 11/74 (14.86%) *Cx. pipiens* (Table 1). Mosquito survival was monitored to assess the impact of viral infection. Control groups fed exclusively on non-infectious blood meals included 82 *Ae. albopictus*, 61 *Ae. aegypti*, and 36 *Cx. pipiens* females. In addition, survival following a second blood feeding was evaluated in *Ae. aegypti* (n = 23) and *Ae. albopictus* (n = 11) females exposed to two consecutive non-infectious blood meals.

**Table 1.** Number and percentage of blood-fed mosquitoes relative to the total number of exposed mosquitoes for each species and experimental replicate. Results are shown separately for 1BF (a single infectious blood meal) and 2BF (a secondary non-infectious blood meal administered after an initial infectious blood meal).

| Species | Replicate | 1BF | 2BF |
| --- | --- | --- | --- |
| *Ae. albopictus* | 1 | 28/92 (30.45%) | - |
|  | 2 | 108/609 (17.73%) | 29/87 (33.33%) |
|  | 3 | 89/343 (25.95%) | 25/47 (53.19%) |
| *Cx. pipiens* | 1 | 12/100 (12%) | - |
|  | 2 | 60/306 (19.61%) | 9/54 (16.67%) |
|  | 3 | 5/97 (5.15%) | - |
|  | 4 | 55/173 (31.79%) | 2/20 (10%) |
|  | 5 | 9/52 (17.31%) | - |
| *Ae. aegypti* | 1 | 72/112 (64.29%) | 14/29 (48.28%) |
|  | 2 | 178/183 (97.27%) | 21/83 (25.30%) |

No significant differences in survival were observed between infected and uninfected mosquitoes in any species (Figure 1A, 1C, 1E). Besides, a second non-infectious blood meal did not affect survival of mosquitoes that had previously ingested one blood meal (Figure 1B, 1D). In contrast, survival differed significantly among species exposed to the infectious blood meal, with *Cx. pipiens* showing lower lifespan than *Aedes* species (Figure 1F; p<0.001).

**Figure 1.**
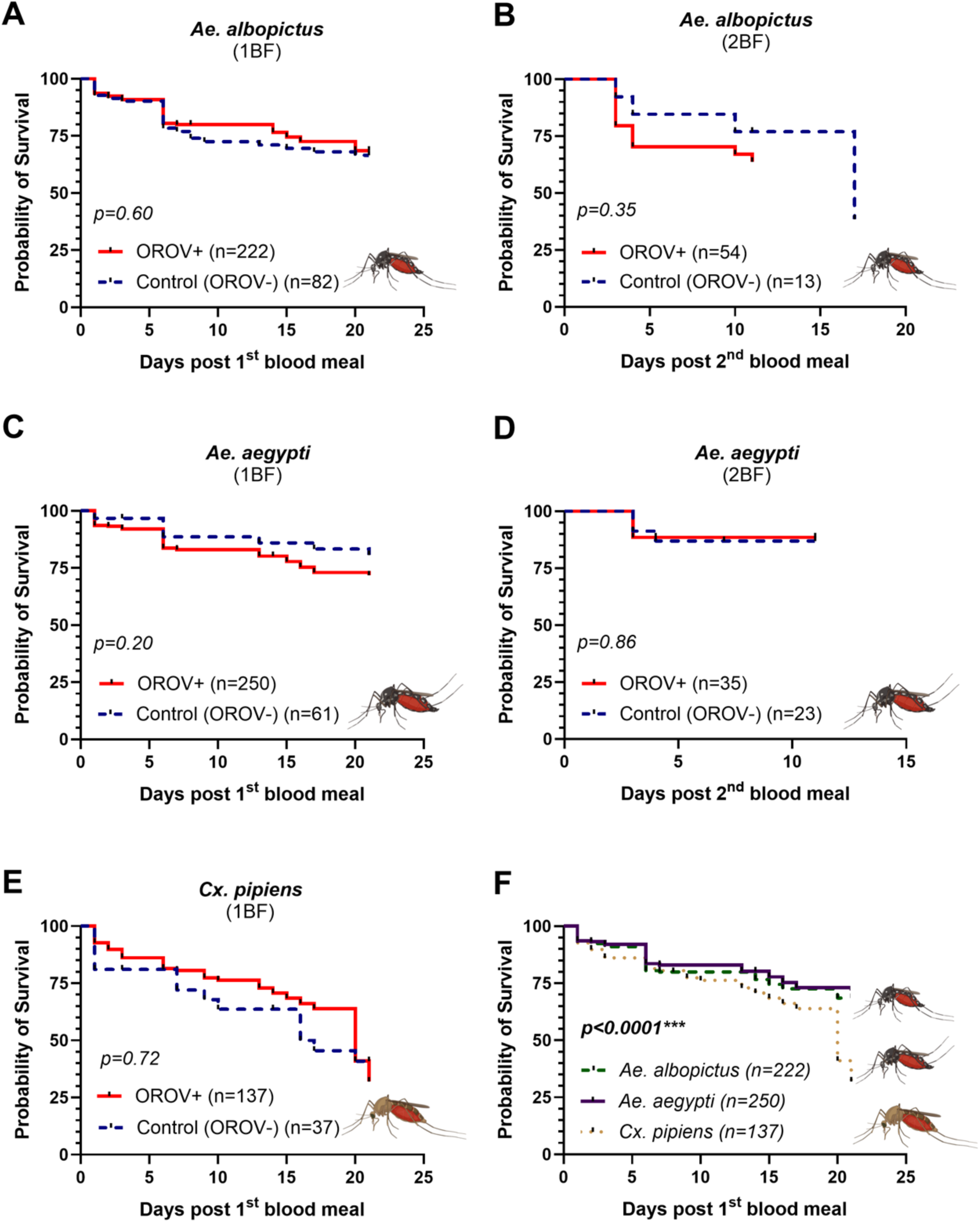
Survival of mosquitoes following Oropouche virus (OROV) exposure. Survival curves of *Aedes aegypti* (A–B), *Aedes albopictus* (C–D) and *Culex pipiens* (E) following exposure to infectious (OROV+) or non-infectious (control, OROV-) bloodmeal with one single infectious blood meal (A, C and E) or with an additional second non-infectious blood meal (B and D)”. (F) Comparative survival curves of the three mosquito species after OROV infectious bloodmeal.

Vector competence was assessed in 159 *Ae. albopictus* (121 from group 1BF and 38 from group 2BF), 160 *Ae. aegypti* (129 from group 1BF and 31 from group 2BF), and 84 *Cx. pipiens* (75 from group 1BF and 9 from group 2BF). Full details on the number of mosquitoes tested per species and sampling time, IR, DR, TR, are presented in Figure 2 and Table 2.

**Figure 2.**
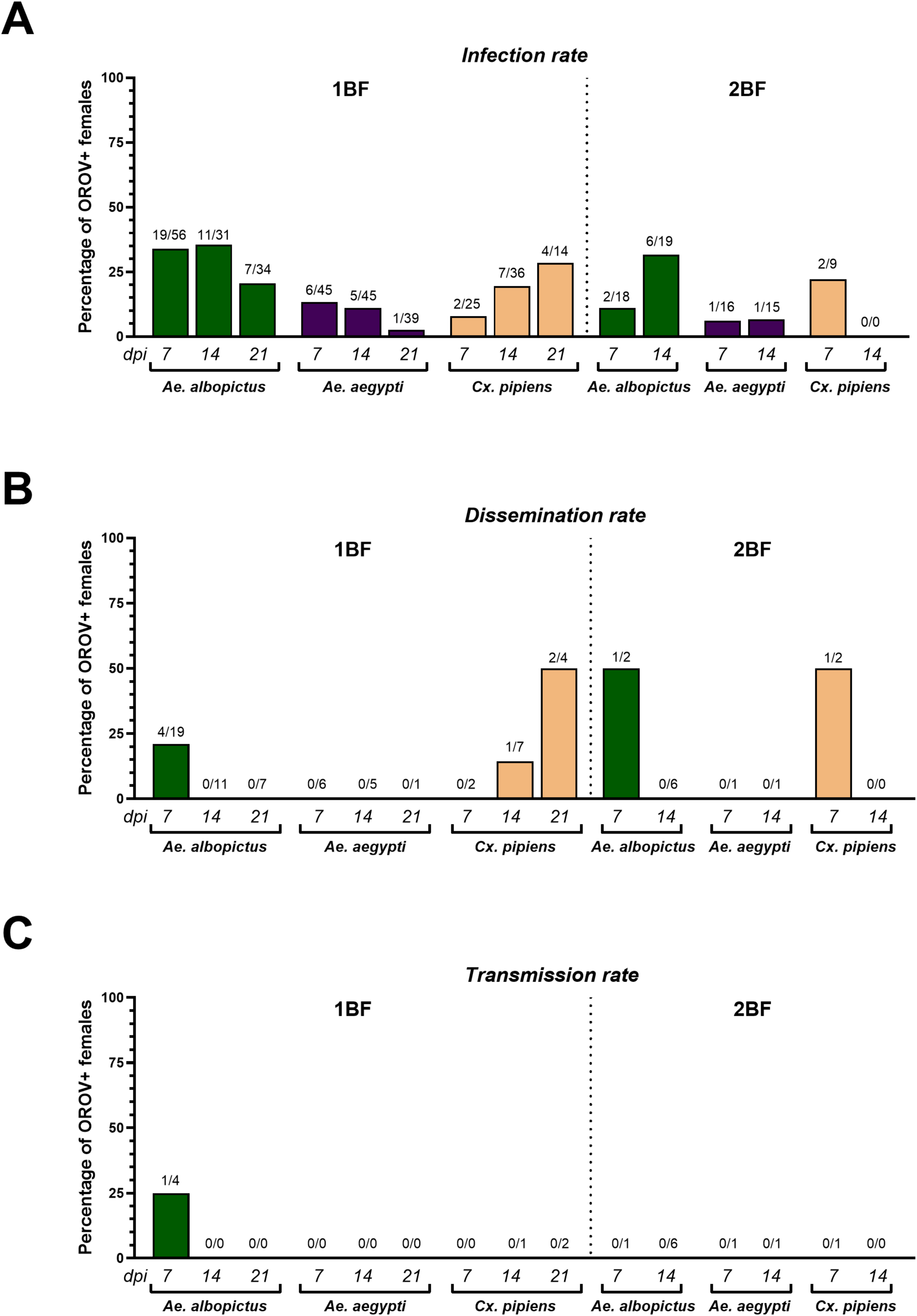
Infection, dissemination and transmission rates of Oropouche virus (OROV) in *Aedes albopictus*, *Aedes aegypti* and *Culex pipiens* following the first (1BF) and second blood feeding (2BF). (A) Infection rates, (B) dissemination rates and (C) transmission rates observed in females analysed at 7-, 14- and 21-days post-infection (dpi) after the first blood feeding, and at 7 and 14 dpi after the second blood feeding. Numbers above bars indicate the number of positive females over the total number tested.

**Table 2.** Infection (IR), dissemination (DR) and transmission rates (TR) for the mosquito species experimentally infected with OROV. Results are shown separately for 1BF (a single infectious blood meal) and 2BF (a secondary non-infectious blood meal administered after an initial infectious blood meal).

| Species | DPI | BF | IR | DR | TR |
| --- | --- | --- | --- | --- | --- |
| *Ae. albopictus* | 7 | 1 | 33.93 (19/56) | 21.05 (4/19) | 25 (1/4) |
|  |  | 2 | 11.11 (2/18) | 50 (1/2) | 0 (0/1) |
| *Ae. albopictus* | 14 | 1 | 35.48 (11/31) | 0 (0/11) | 0 |
|  |  | 2 | 31.58 (6/19) | 0 (0/6) | 0 |
| *Ae. albopictus* | 21 | 1 | 20.59 (7/34) | 0 (0/7) | 0 |
|  |  | 2 | - | - | - |
| *Cx. pipiens* | 7 | 1 | 8 (2/25) | 0 (0/2) | 0 |
|  |  | 2 | 22.22 (2/9) | 50 (1/2) | 0 (0/1) |
| *Cx. pipiens* | 14 | 1 | 19.44 (7/36) | 14.29 (1/7) | 0 (0/1) |
|  |  | 2 | - | - | - |
| *Cx. pipiens* | 21 | 1 | 28.57 (4/14) | 50 (2/4) | 0 (0/2) |
|  |  | 2 | - | - | - |
| *Ae. aegypti* | 7 | 1 | 13.33 (6/45) | 0 (0/6) | 0 |
|  |  | 2 | 6.25 (1/16) | 0 (0/1) | 0 |
| *Ae. aegypti* | 14 | 1 | 11.11 (5/45) | 0 (0/5) | 0 |
|  |  | 2 | 6.66 (1/15) | 0 (0/1) | 0 |
| *Ae. aegypti* | 21 | 1 | 2.56 (1/39) | 0 (0/1) | 0 |
|  |  | 2 | - | - | - |

DPI, Days post-infection
BF, Blood-feeding group

For mosquitoes of the 1BF group, *Ae. albopictus* exhibited IR of 33.93% at 7 dpi, 35.48% at 14 dpi, and 20.59% at 21 dpi. Dissemination occurred in 21.05% of infected individuals at 7 dpi, but it was absent at 14- and 21 dpi. Transmission was detected only once (1/4) at 7 dpi (Table 2, Figure 2), corresponding to the individual that exhibited the highest viral load in both the body and legs (Figure 3). Mean viral titres in this individual were of 3.44×10^2^ TCD₅₀/ml in the body, 3.2×10^2^ TCD₅₀/ml in the legs, and 2.9×10⁻³ TCD₅₀/ml in the saliva (Figure 3). Despite inoculation of one third of the saliva sample of that mosquito (100 μl) on Vero cells, no cytopathic effect (CPE) was observed after three passages. In the 2BF group, IR were 11.11% at 7 dpi and 31.58% at 14 dpi. Dissemination was detected in 50% of infected individuals at 7 dpi but not at 14 dpi, and no transmission was observed (Figure 2).

**Figure 3.**
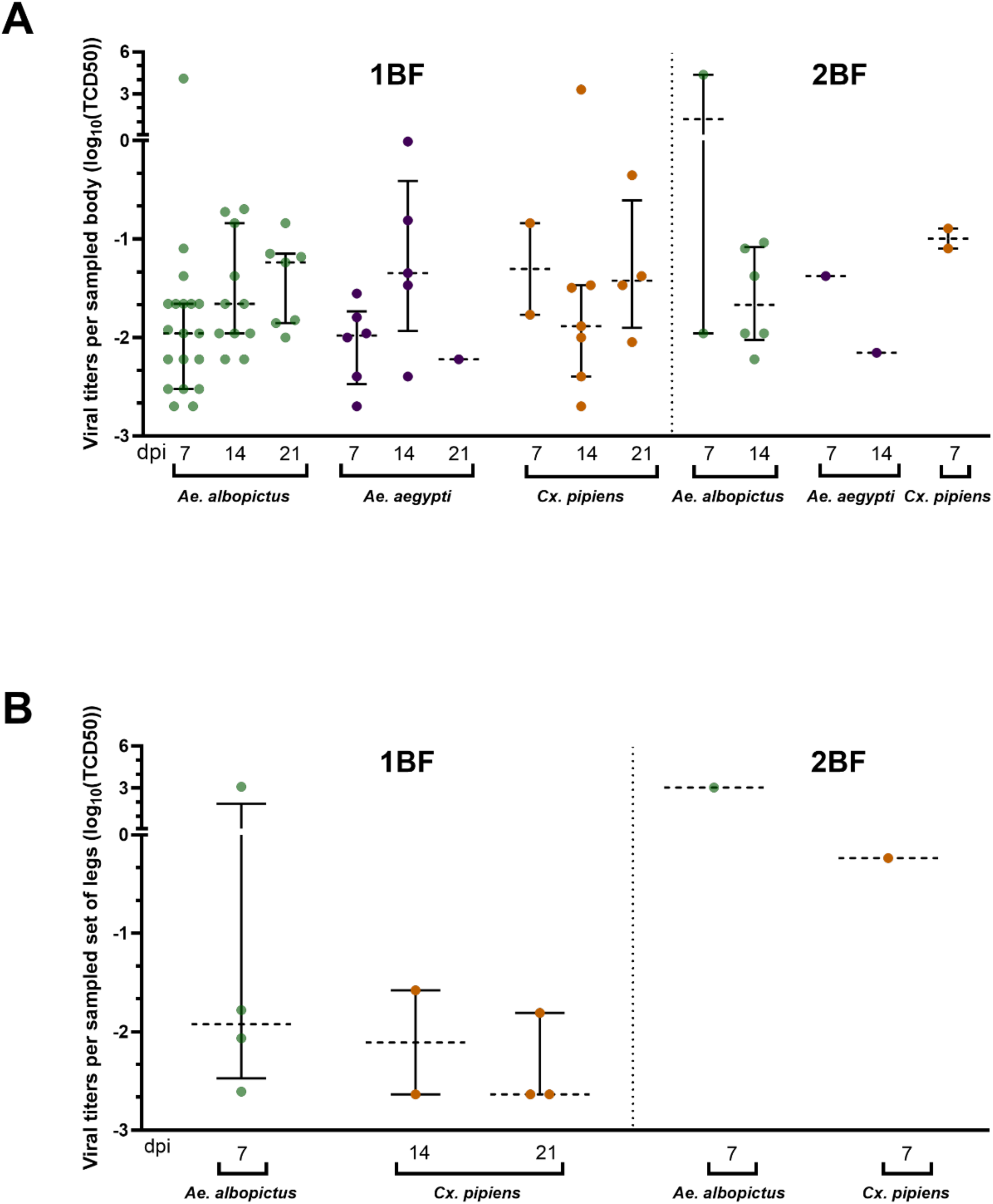
OROV viral titer per sample. Individual points represent obtained viral titer of bodies (A) and legs (B) of OROV-positive individuals. Data were grouped by mosquito species, day post-infection (dpi), and feeding status (1BF, a single infectious blood meal; 2BF, a secondary non-infectious blood meal given to mosquitoes after an initial infectious blood meal). Median and interquartile values are shown for each study group.

For *Ae. aegypti*, in the 1BF group, IR were 13.33% at 7 dpi, 11.11% at 14 dpi, and 7.69% at 21 dpi. No dissemination or transmission was observed in *Ae. aegypti*. In the 2BF group, IR were 6.25% at 7 dpi and 6.67% at 14 dpi, with no events of dissemination or transmission (Table 2, Figure 2).

For *Cx. pipiens*, in the 1BF group, IR increased from 8% at 7 dpi to 19.44% at 14 dpi, and 28.57% at 21 dpi. Dissemination was observed in 14.29% of infected mosquitoes at 14 dpi and 50% at 21 dpi, and transmission was never detected. In the 2BF group, 22.22% of mosquitoes became infected at 7 dpi, with dissemination in one of them (50%). Transmission was absent at all time points (Table 2, Figure 2).

IR in mosquito bodies did not differ across species (χ² = 4.54, df = 2, p = 0.10), days post-infection (χ² = 2.07, df = 1, p = 0.15), or blood-feeding treatments (1BF vs. 2BF; χ² = 2.26, df = 1, p = 0.13). Likewise, the interaction between species and blood-feeding group was not significant (χ² = 0.47, df = 2, p = 0.78). Regarding DR, we did not detect significant effects of species (χ² = 2.69, df = 2, p = 0.26), blood-feeding treatment (1BF vs. 2BF; χ² = 0.14, df = 1, p = 0.71), or days post-infection (χ² = 1.33, df = 1, p = 0.25).

TR was detected in only one *Ae. albopictus* mosquito, preventing any meaningful statistical analysis of TR across species, feeding groups, or days post-infection.

The viral concentrations in mosquito bodies did not differ significantly between species (χ² = 1.43, df = 2, p = 0.49), days post-infection (χ² = 0.98, df = 1, p = 0.32), or blood-feeding treatments (χ² = 2.99, df = 1, p = 0.08). Likewise, the interaction between species and blood-feeding group was not significant (χ² = 2.28, df = 2, p = 0.32). The viral concentrations in mosquito legs did not differ significantly between species (χ² = 3.78, df = 1, p = 0.052), blood-feeding groups (χ² = 2.85, df = 1, p = 0.09), or sampling times (χ² = 1.04, df = 1, p = 0.31).

Finally, across all three mosquito species, no viral RNA was detected in the F1 adult pools analyzed, suggesting that vertical transmission did not occur in this study.

## Discussion

In this work, we evaluated the vector competence of the most common mosquito species present in Spain, including Spanish populations of *Ae. albopictus* and *Cx. pipiens,* for the emergent OROV strain responsible for the 2024 outbreak. Additionally, given the major role of *Ae. aegypti* as a global arbovirus vector and its recent detection in the Canary Islands [16, 17], *Ae. aegypti* (Liverpool strain) was also included in the assays. We also tested the effect of a second blood meal on vector competence, as well as the potential vertical transmission of OROV. Our results show that *Ae. aegypti* and *Cx. pipiens* are not competent vectors for this OROV strain, while *Ae. albopictus* exhibits limited susceptibility. Although OROV RNA was detected in the saliva of a single *Ae. albopictus* at 7 dpi, the low viral load, the absence of CPE in cell culture, and the unsuccessful virus isolation after three passages on VERO cells, suggest that this finding could reflect residual RNA rather than biologically meaningful transmission.

The epidemiological relevance of these findings is underscored by the widespread distribution of some of these mosquito species across Europe [17, 30], and particularly in Spain, which has reported the highest number of imported OROV cases in Europe to date [3], increasing the probability of virus–vector encounters in areas where competent or partially competent mosquito species are established. *Aedes albopictus* and *Cx. pipiens* dominate entomological surveys in urban areas of mainland Spain and the Balearic Islands [30, 31], while *Ae. aegypti* has been recently detected in the Canary Islands [17], and is established in other European regions, including Madeira [32] and parts of the Balkans [16]. Under these circumstances, experimental assessments of vector competence using local mosquito populations are essential to accurately evaluate the risk of autochthonous transmission and to inform surveillance and preparedness strategies.

Given the interest in Europe in evaluating the risk of autochthonous transmission of OROV, European groups have evaluated the vector competence of mosquito species such as *Ae. albopictus* or *Cx. pipiens* for OROV, reporting variable and sometimes contradictory outcomes, ranging from a complete absence of transmission to successful viral dissemination and, in some cases, transmission [13–15]. In Europe, recent studies using *Ae. albopictus* populations have generally reported low or limited infection and transmission rates for OROV. For example, Jansen *et al.* [14] observed low infection rates that varied with temperature and time post-infection, with OROV strain TR 9760 (1955, Trinidad and Tobago) transmission detected only at 14 dpi in mosquitoes reared at 24 °C and at 21 dpi at 27 °C. Rosales-Rosa *et al.* [15] also documented low infection rates using the same strain (TRVL9760 strain; 1955, Trinidad and Tobago; accession numbers KC759122–24) and a strain isolated in 2024 from a patient of Cuba (OROV-IRCCS-SCDC_1/2024, imported from Cuba to Italy; GenBank accession numbers PP952117–19), with viral dissemination detected in infected mosquitoes but no virus detected in saliva. Similarly, Mancuso *et al.* [13] detected viral RNA in the bodies of a small number of individuals from an Italian *Ae. albopictus* population, but no dissemination or transmission was observed using also the strain OROV-IRCCS-SCDC_1/2024. Overall, these studies indicate that European *Ae. albopictus* populations display low susceptibility to OROV, with infection occurring at low rates and transmission being detected only under restricted experimental conditions.

The absence of OROV transmission observed in *Cx. pipiens* in our study is consistent with patterns reported in previous experimental studies. In *Cx. pipiens* populations from Italy, no infection, dissemination or transmission was detected [13]. Similarly, Rosales-Rosa *et al.* [15] reported that *Cx. pipiens* from Belgium was not susceptible and did not support viral replication leading to transmission. Although infection was detected in a *Cx. pipiens* population from Germany, no virus was found in saliva, in agreement with our results [14]. To date, only a single study conducted in the United States has reported OROV transmission by *Cx. pipiens*, with viral RNA detected in the saliva of one individual using the strain OROV 240023 (GenBank accession number PQ064919.1, PQ064920.1, and PQ064921.1) [34]. Overall, OROV competence in *Cx. pipiens* appears to be limited and may vary geographically or depend on specific ecological or virological factors. These isolated observations highlight the need for further intraspecific assessments across diverse *Cx. pipiens* populations.

Despite being a globally important arbovirus vector, competent for dengue virus (DENV), CHIKV, and ZIKV [35], *Ae. aegypti* LVP strain, showed limited dissemination of OROV in our study.

Similarly, previous studies also reported only transient infection and low IR, but no transmission, indicating that, *Ae. aegypti* is unlikely to support efficient OROV transmission [11, 35].

Overall, evidence gathered across multiple studies from different research groups, including our work, indicates that OROV shows a limited capacity to infect, disseminate and be transmitted by major European mosquito species, such as *Cx. pipiens* and *Ae. albopictus*. Nevertheless, contrasting results reported among European studies highlight the complexity of OROV-mosquito interactions and suggest that vector competence may be influenced by multiple factors, including viral strain, mosquito population genetics, temperature, and experimental design [36]. Several previous studies have relied on limited sample sizes or assessed vector competence at only one or two time points during the extrinsic incubation period. As a result, additional experimental studies are needed to better characterize the kinetics of OROV replication in mosquitoes, ideally incorporating multiple incubation temperatures, different viral strains, and larger experimental cohorts, to more accurately assess the potential for OROV emergence in Europe. In this context, methodological heterogeneity across studies-including differences in infectious dose, virus strain and incubation temperature-likely contributes to the variability reported [36, 37]. These limitations underscore the need for more standardized experimental approaches [38], to facilitate robust comparisons and improve risk assessment of emerging arboviruses.

Our findings suggest that the principal barriers to OROV infection and transmission are likely located at the midgut level rather than resulting from intrinsic molecular incompatibilities between the virus and its mosquito hosts. Although viral dissemination to secondary tissues, as evidenced by virus detection in legs, was occasionally observed, these events were rare and occurred at very low frequencies. This pattern indicates that OROV infection is largely restricted at the level of midgut infection and/or midgut escape, thereby limiting subsequent viral dissemination and transmission [39].

Numerous studies have demonstrated that a second non-infectious blood meal significantly shorten the extrinsic incubation period of arboviruses such as DENV, CHIKV or ZIKV in mosquitoes (e.g., *Ae. aegypti* and *Ae. albopictus*), by enhancing viral dissemination from the mosquito midgut, possibly through microperforations in the basal lamina [22, 23]. However, we did not observe significant differences in OROV dissemination between individuals receiving a second blood meal and those fed only once, in any of the species analyzed which agrees with recent published data where multiple blood meals did not affect OROV dissemination in *Ae. aegypti* [24]. Similarly, other studies, using the same *Ae. albopictus* strain than in our study, have reported no effect of a second non-infectious blood meal on vector competence for other arbovirus [40]. However, the small sample sizes, particularly the limited number of *Cx. pipiens* in the double-feeding group, may have reduced our statistical power to detect effects of secondary blood meals on OROV dissemination.

The absence of virus particles in all F1 adult pools from the species analyzed indicates that vertical transmission did not occur under the conditions tested, as also reported in previous studies [13]. However, as highlighted by these authors, results obtained from a single gonotrophic cycle may not be fully representative of the potential for viral transmission to the eggs. Therefore, the lack of viral detection in the F1 generation cannot definitively rule out the occurrence of transovarial transmission. Additional studies incorporating multiple gonotrophic cycles would be required to robustly assess the likelihood of vertical transmission, particularly under scenarios in which these mosquito populations might become competent vectors.

## Conclusion

Our results indicate a limited vector competence of Spanish *Ae. albopictus* and a lack of competence of Spanish *Cx. pipiens* and the *Ae. aegypti* LVP strain for the emergent OROV strain. Overall, these findings suggest a low potential for mosquito-borne OROV transmission under current conditions in Europe. Nevertheless, co-evolutionary processes and shifts in vector–virus interactions that could facilitate OROV adaptation to new environments should not be overlooked. The recent detection of imported OROV fever cases in Spain and other European countries underscores the need for continuous vector surveillance and further research on the ecological and climatic factors that may influence OROV emergence, including experimental evaluation of *Culicoides* populations to assess the risk of local transmission.

## Acknowledgments

Thanks to F.N.G. for his help in the laboratory. We thank to the biosafety department of the National Center for Microbiology for their support. We also acknowledge the Instituto de Salud Carlos III (ISCIII) for the financial support provided for the maintenance and operation of the insectary facilities.

## Author contribution

**R.G.L. and I.M.M.** conceived the study. **R.G.L., I.M.M., and N.L.** performed experiments. **R.G.L., I.M.M., and M.L.d.F.** curated and analyzed the data. **R.G.L.** wrote the original draft. **R.G.L., M.S.-S., M.J., I.M.M. and A.V.** acquired funding. **P.S.-M., M.L.d.F.**, **S.D.-E., A.M.-R., A.P., E.P.-M., R.M., M.S.-S., M.J., I.M.M, and A.V.** contributed to writing, reviewed and edited the manuscript, and approved the final version.

## Funding

This work was supported by CIBER (Consorcio Centro de Investigación Biomédica en Red) (group CB21/13/00110), Instituto de Salud Carlos III, Ministerio de Ciencia e Innovación and Unión Europea-NextGenerationEU. This research was also supported by project INFEC24PI05 S.N., funded by the strategic action of the CIBER of Infectious Diseases (CIBERINFEC) 2024, the Intramural Call for Research Projects CIBERINFEC-2024. RGL is currently funded by a Sara Borrel postdoctoral contract (Ref. CD22CIII-00009) from the Instituto de Salud Carlos III. MdL is funded by a FPU, “Formación del Profesorado Universitario”, grant from the Ministry of Education of the Government of Spain (Ref. FPU23/01929). PSM is funded by a PFIS contract (Ref. FI22CIII/00040) from the Instituto de Salud Carlos III.

## Conflict of interest

The authors declare no conflict of interest.

## Notes

### Competing Interest Statement

The authors have declared no competing interest.

